# S100A9 Inhibits and Redirects Prion Protein 89-230 Fragment Amyloid Aggregation

**DOI:** 10.1101/2024.02.06.579161

**Authors:** Mantas Ziaunys, Darius Sulskis, Kamile Mikalauskaite, Andrius Sakalauskas, Ruta Snieckute, Vytautas Smirnovas

## Abstract

Protein aggregation in the form of amyloid fibrils has long been associated with the onset and development of various amyloidoses, including Alzheimer’s, Parkinson’s or prion diseases. Recent studies of their fibril formation process have revealed that amyloidogenic protein cross-interactions may impact aggregation pathways and kinetic parameters, as well as the structure of the resulting aggregates. Despite a growing number of reports exploring this type of interaction, they only cover just a small number of possible amyloidogenic protein pairings. One such pair is between two neurodegeneration-associated proteins: the pro-inflammatory S100A9 and prion protein, which are known to co-localize *in vivo*. In this study, we examined their cross-interaction *in vitro* and discovered that the fibrillar form of S100A9 modulated the aggregation pathway of mouse prion protein 89-230 fragment, while non-aggregated S100A9 also significantly inhibited its primary nucleation process. These results complement previous observations of the pro-inflammatory protein’s role in amyloid aggregation and highlight its potential role against neurodegenerative disorders.

## Introduction

Amyloid fibril formation is a key factor linked with the onset and progression of amyloidoses, including neurodegenerative Alzheimer’s, Parkinson’s or prion diseases ^1,2^. Currently, multiple proteins and peptides of various lengths are known to aggregate into such highly organized, beta-sheet rich structures ^1^. In the last decade, due to the advancement of structural biology within the cryogenic electron microscopy (Cryo-EM) field, numerous structures of amyloid fibrils were solved ^3^. Consequently, this led to the detailed findings that the same proteins form a multitude of different fibrils, depending strongly on the environment and aggregation conditions ^4,5^. The complex nature of amyloid formation is one of the reasons why there are still very few disease modulating drugs available, with many promising candidates that show efficacy during initial studies often failing to pass clinical trials ^6,7^. Considering that the sum of amyloid-related disorders is in the tens of millions and the number of affected individuals is predicted to increase even further ^8,9^, it is essential to decipher this process in the hopes of discovering a potential cure.

The fundamental model of amyloid aggregation ^10^ consists of nucleation ^11,12^, elongation ^13,14^ and two secondary processes (fibril fragmentation ^15^ and surface-induced nucleation ^16,17^). However, this model primarily describes single protein reactions, while the actual cell environment is densely crowded with numerous proteins and thus the aggregation mechanisms become affected by them ^18^. Currently, there are multiple reports which show that amyloidogenic proteins are capable of interacting with one another, altering their aggregation pathways and resulting fibril structural characteristics. For instance, Alzheimer’s disease-related amyloid beta peptide was able to influence the aggregation of multiple neurodegenerative disease-related proteins, including alpha-synuclein (Parkinson’s disease) ^19^, Tau (Alzheimer’s disease) ^20,21^, and prion proteins (prion diseases) ^22^. Similarly, alpha-synuclein interacted with Tau ^23,24^ and prion proteins ^25^. In total, according to the amyloid-amyloid interactions database ^26^ (2023), amyloid-beta peptide interacts with 22 and alpha-synuclein with 17 unique proteins with more discovered each year Despite a growing number of such observations, the overall picture of the amyloid interactome remains incomplete.

One potential cross-interaction, which has received minimal attention, is between S100A9 and prion protein. S100A9 is part of a pro-inflammatory S100 calcium-binding protein family ^27^, members of which have been implicated in Alzheimer’s disease, either through their association with amyloid plaques, effects on signalling pathways or interactions with amyloid beta ^28^. Other studies have demonstrated their involvement with alpha-synuclein aggregation ^29^, as well as Tau protein hyperphosphorylation ^30^, suggesting that they may play an important role in the onset of amyloid disorders. Interestingly, S100A9 has also been shown to be capable of rapidly forming amyloid-like structures in apo-form under physiological conditions without any lag phase ^31^.

Prion proteins, on the other hand, are cell-surface glycoproteins, which perform multiple physiological functions, such as copper homeostasis, neuroprotection and stem cell renewal ^32^. Their amyloid aggregate formation and accumulation is associated with several disorders, including Creutzfeldt-Jakob disease, Gerstmann–Straussler–Scheinker syndrome, fatal familial insomnia ^33,34^ and spongiform encephalopathy ^35^. These numerous disease phenotypes stem from prion protein ability to form a large variety of aggregate types. Such different fibril strains can be the result of multiple factors, including point mutations, cross-interactions with other proteins or the environmental conditions under which the aggregation occurs ^36–38^. Despite much progress in regards to prion protein aggregate polymorphism and their relation to neurodegenerative disorders, there is still no definite explanation for their structure-condition relationship.

According to the PAXdb Protein abundance database ^39^, S100A9 and prion proteins share a localization in multiple areas *in vivo*, including the cerebral cortex ^40,41^ and cerebrospinal fluid ^42^. These proteins also play roles in neuroinflammation, either by pro-inflammatory signaling of native S100A9, cytotoxicity of fibrillar S100A9 ^28^ or activation and proliferation of microglia by prion protein aggregates ^43^. Additionally, both S100A9 and prion proteins are capable of binding metal ions, including Ca^2+^ and Zn^2+^, which have been shown to modulate their oligomerization and amyloid aggregation. For example, Ca^2+^ ions inhibit S100A9 aggregation ^44^, while at the same time promoting the cytotoxic structure of prion protein ^45^. These factors, coupled with existing reports of S100A9 and prion protein cross-interactions with other amyloidogenic proteins ^28,46,47^, prompt the need for more in-depth investigations into this pairing.

In this work, we investigated the cross-interaction between S100A9 and a mouse prion protein 89-230 fragment, which comprises the proteinase-K resistant region found in infectious forms of the protein ^48^. We discovered that the fibrillar form of S100A9 modulated the prion protein fragment’s aggregation pathway, while the non-aggregated form also strongly inhibited the primary nucleation process. These results, along with previous studies of S100A9 cross-interaction with amyloidogenic proteins, indicate that the pro-inflammatory S100A9 may serve a potential role in combating the onset and progression of neurodegenerative amyloid disorders.

## Materials and Methods

### Protein aggregation assays

Mouse prion protein fragment 89-230 (further referred to as PrP) was purified as described previously ^49^, without the His-tag cleavage step, dialysed in 10 mM sodium acetate buffer solution for 24 hours, concentrated to 200 μM and stored at −20°C. S100A9 was purified as described in Supplementary Materials, concentrated to 300 μM in PBS buffer solution (pH 7.4, 137 mM NaCl, 2.7 mM KCl, 10 mM Na_2_HPO_4_, 1.8 mM KH_2_PO_4_) and stored at −20°C. For all further experimental procedures, the same batches of prion protein and S100A9 were used in order to avoid possible batch-to-batch variability between different purifications.

Thioflavin-T (ThT) was dissolved in Milli-Q H_2_O (∼11 mM concentration) and filtered through a 0.22 μm pore size syringe filter. ThT concentration was measured by scanning a diluted dye sample’s absorbance at 412 nm, using a Shimadzu UV-1800 spectrophotometer (ε_412_=23250 M^−1^cm^−1^). The dye was then diluted to 10 mM and stored at −20°C under dark conditions. To account for a higher dye binding propensity of fibrils under high ionic strength conditions ^50^, as well as ThT hydroxylation at pH 7.4 ^51^, a dye concentration of 100 μM was chosen for all further kinetic monitoring procedures. Based on a fluorescence assay, larger dye concentrations would lead to high levels of self-quenching and other fluorescence-distorting effects (Supplementary Figure S1).

To monitor PrP aggregation kinetics under various guanidine hydrochloride (GuHCl) concentrations, the PrP solution was combined with 6 M GuHCl PBS, 1x PBS, 10x PBS and 10 mM ThT solutions to yield a final protein concentration of 25 μM, 100 μM ThT, 1x PBS and 0 – 3.0 M GuHCl. The reaction solutions were then distributed to 96-well non-binding plates (cat. No 3881, Fisher Scientific, each well contained 100 μL solution and a 3 mm glass bead, 6 wells for each condition), sealed by Nunc-sealing tape and incubated at 37°C with constant 600 RPM agitation in a ClarioStar Plus plate reader. ThT fluorescence was monitored every 5 minutes using 440 nm excitation and 480 emission wavelengths.

To prepare S100A9 fibrils and monitor their formation kinetics, the S100A9 stock solution was combined with 6 M GuHCl PBS, 1x PBS and 10 mM ThT to a final protein concentration of 200 μM, 100 μM ThT and 0 or 1.5 M GuHCl. Samples, which were used to determine the residual non-aggregated S100A9 concentration at the end of the reaction, did not contain ThT (due to ThT absorbance at 280 nm). The aggregation reaction and ThT fluorescence monitoring was then conducted as in the case of PrP. For further use of S100A9 aggregates, the solutions from each well were recovered and combined. To determine the concentration of non-aggregated S100A9 in these samples, they were centrifuged at 14 000 x g for 15 minutes and the absorbance of the supernatant was measured using a Shimadzu UV-1800 spectrophotometer (ε_280_ = 7100 M^−1^cm^−1^).

To monitor PrP aggregation in the presence of S100A9, the stock PrP solution was combined with either non-aggregated S100A9 or the previously prepared fibril samples, 6 M GuHCl PBS, 1x PBS, 10x PBS and 10 mM ThT. The resulting solutions contained 25 μM PrP, 100 μM ThT, 1.5 M GuHCl, 1x PBS and a range of S100A9 concentrations (0, 1, 2.5, 5, 25 and 50 μM). The aggregation reaction and ThT fluorescence monitoring was then conducted as in both other previously described cases (16 wells for each condition). To examine the influence of Ca^2+^ on the aggregation kinetics, the reaction solutions were additionally combined with PBS containing 5 mM CaCl_2_, to a final Ca^2+^ concentration of 0.5 mM (10-times the concentration of S100A9).

All reaction lag time and apparent rate constant values were determined as described previously ^46^. In brief, each sigmoidal curve was fit using a Boltzmann sigmoidal equation and lag time, apparent rate constants were determined from the fit parameters. For seeded aggregation, the reaction t_50_ values were determined by normalizing the curves between 0 and 1 and applying a linear fit between 0.4 and 0.6 intensity value data points. The time point at 0.5 value of the linear fit Y-axis was regarded as the reaction’s t_50_ value. All data analysis was done using Origin software.

### Prion protein self-replication

To prepare PrP samples for Fourier-transform infrared spectroscopy, aliquots of each PrP aggregate sample (40 μL) were combined with the initial reaction solution (360 μL, which did not contain S100A9) and incubated under identical conditions as were used in the initial kinetic monitoring assay (4 wells for each condition). This procedure was repeated twice to dilute the concentration of S100A9 in the solution 100 times (10-time dilution of initial sample S100A9 concentration with each round of reseeding). The self-replication reactions were conducted until the kinetic curves reached a plateau, indicating the conclusion of the process (total incubation time – 16 hours).

To measure PrP self-replication in the presence of S100A9, 60 μL aliquots of PrP fibrils (prepared in the absence of S100A9) were combined with the initial reaction solutions (540 μL), which were supplemented with either non-aggregated S100A9 or S100A9 fibrils. The final reaction solutions contained 25 μM PrP, 0 – 50 μM S100A9, 100 μM ThT, 1.5 M GuHCl and 1x PBS. Aggregation monitoring was done as in previous cases (6 wells for each condition).

### Aggregate stability against denaturation

The prepared S100A9 aggregate samples were combined with 1x PBS and 6 M GuHCl PBS, resulting in solutions with 40 μM S100A9 and a range of GuHCl concentrations (0 – 4.5 M with 0.1 M increments). The solutions were then distributed to 200 μL test-tubes and incubated at 37°C for 24 hours without agitation. After incubation, they were distributed to 96-well non-binding plates (100 μL volume in each well) and their optical densities at 600 nm (OD_600_) were scanned using a ClarioStar Plus plate reader (3 technical repeats). The optical density values were corrected by subtracting the OD_600_ values of control solutions, which did not contain S100A9.

### Fourier-transform infrared spectroscopy (FTIR)

200 μL aliquots of each sample were centrifuged at 14 000 x g for 15 minutes, after which the supernatant was carefully removed and replaced with 200 μL of D_2_O, containing 400 mM NaCl ^50^. The centrifugation and resuspension procedure was repeated 4 times. The resulting sample FTIR spectra were acquired as previously described ^52^, using a Bruker Invenio S FTIR spectrometer. For each sample, 256 spectra were acquired and averaged. The spectra were then baseline corrected and normalised to the same band area in the range between 1595 cm^−1^ and 1700 cm^−1^. Spectra deconvolution was done as described previously ^52^, using a Gaussian peak-fitting function, number of peaks – 4, RMS noise – 0.04. Data processing was done using GRAMS software.

PrP and S100A9 spectra were deconvoluted into PrP Type I, PrP Type II and S100A9 fibril spectra by applying the least squares mathematical method. For this procedure, the normalized spectra of Type I, Type II PrP and S100A9 fibrils were combined using an array of different percentages of each spectrum (0 – 100% at 0.01% increments). The resulting combined spectrum was then compared to the normalized experimental result spectrum. The combined spectrum with the lowest deviation was determined using the least squares method and served as a basis for calculating the sample composition. For this calculation, the number of amide bonds contributing to the total FTIR Amide I band spectra by PrP was 163, S100A9 – 113.

### Atomic force microscopy

40 μL aliquots of samples were placed on freshly cleaved mica, incubated for 2 minutes at room temperature, washed with 2 mL of Milli-Q H_2_O and dried using airflow. The AFM image acquisition was done as described previously ^52^, using a Bruker Dimension Icon atomic force microscope. 1024 x 1024 pixel resolution images were flattened using Gwyddion 2.57 software ^53^. Fibril cross-sectional heights were determined by tracing lines perpendicular to the fibril axes (n=30).

### Protein melt assay

8-Anilinonaphthalene-1-sulfonic acid (ANS) was dissolved in Milli-Q H_2_O (∼6 mM concentration) under dark conditions and filtered through a 0.22 μm pore size syringe filter. ANS concentration was measured by scanning a diluted dye sample’s absorbance at 351 nm, using a Shimadzu UV-1800 spectrophotometer (ε_351_=5500 M^−1^cm^−1^). The dye was diluted to 5 mM and used immediately after preparation. PrP and S100A9 stock solutions were combined with 1x PBS and 5 mM ANS to final protein concentrations of 100 μM and 100 μM ANS. The samples were then placed in 3 mm pathlength quartz cuvettes (200 μL volume, sealed with a plug cap) and ANS fluorescence was measured using a Varian Cary Eclipse spectrofluorometer. The protein melt assay was conducted by incubating the samples at 25°C for 5 minutes, followed by a gradual temperature increase of 2°C/min. Measurements were taken every 30 seconds, using 370 nm excitation and 470 emission wavelengths, 2.5 s signal averaging time. For each condition, the protein melt assay was repeated 3 times. The protein melting point temperature (T_m_) values were compared by calculating the first derivative of the fluorescence curves (10 point Savitzky-Golay smoothing) and determining their maximum value positions.

### Size-exclusion chromatography

Size exclusion chromatography (SEC) was performed with a Tricorn 10/300 column loaded with Superdex 75 resin (Cytiva) on ÄKTA start protein purification system (Cytiva) at 4 °C. Prior to sample loading, the column was equilibrated with PBS buffer, supplemented with 1.5 M GuHCl. Approximately 250 μL of sample (prepared in the same buffer solution) containing either a single native protein solution or a mixture of both non-aggregated S100A9 (100 μM) and PrP (100 μM) were injected into a column.

### MTT and LDH assays

Aggregated PrP samples were centrifuged at 14 000 x g for 15 minutes, after which the supernatants were carefully removed, replaced with an identical volume of PBS solution. The centrifugation and resuspension procedure was repeated 3 times to remove all residual denaturant. After the final centrifugation, the aggregate pellet was resuspended into a 2-times smaller volume. The MTT and LDH assays were conducted as described previously ^54^. For both assays, the final concentration of aggregated PrP was 10 μM (assuming complete conversion of monomeric PrP into aggregates). For each condition, 3 independent assays were conducted, each with 3 technical repeats.

## Results

In order to examine how both the non-aggregated and amyloid fibril forms of S100A9 influence the aggregation process of mouse prion protein 89-230 (PrP), it was first necessary to determine suitable experimental conditions. In an ideal case, meeting three key criteria was essential for obtaining reliable results: a) the prion protein aggregation reaction concludes in a reasonable timeframe (to avoid gradual sample evaporation and fluorescent dye hydroxylation ^51^), b) the generated S100A9 fibrils remain stable under the selected conditions, c) the spontaneous aggregation of S100A9 is significantly hindered. Since prion protein aggregation *in vitro* is often conducted under denaturing conditions, adjustment of this factor was chosen as a basis for achieving all three criteria.

PrP aggregation under a range of guanidine hydrochloride (GuHCl) concentrations (Figure 1A) revealed that the shortest lag time for PrP fibril formation was around 2.2 M (average value with one standard deviation was 124 ± 11 minutes). Both lower and higher denaturant concentrations resulted in substantially longer lag time periods (1455 ± 159 minutes at 1 M GuHCl and 307 ± 28 minutes at 3 M GuHCl). In the case of 1.0 – 1.2 M GuHCl, the aggregation was also significantly more stochastic with larger deviations in lag time values. Based on these results, suitable GuHCl concentrations for PrP – S100A9 coaggregation studies would range from 1.4 M to 3.0 M.

**Figure 1.**
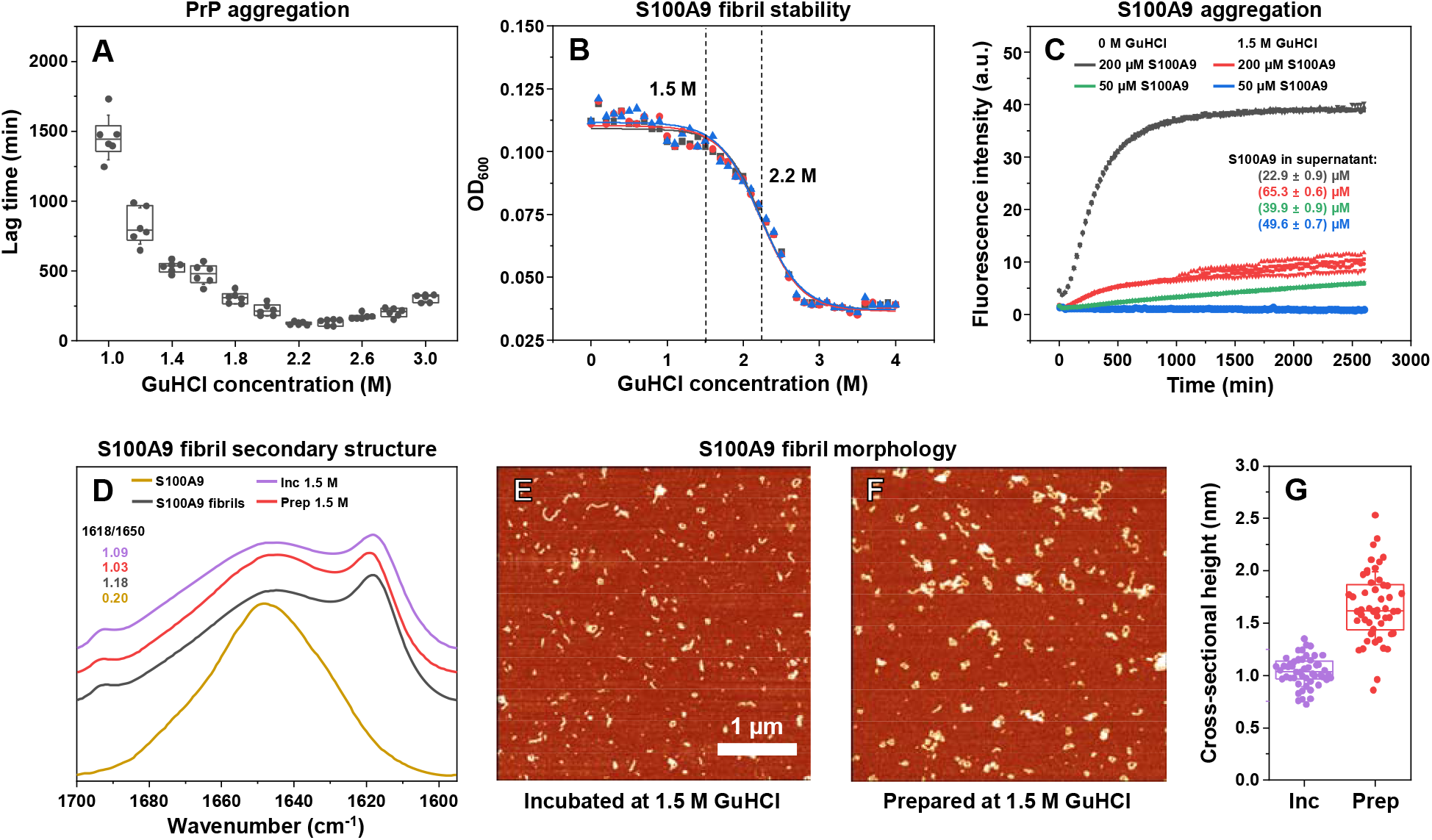
Influence of guanidinium hydrochloride (GuHCl) concentration on prion protein and S100A9 aggregate formation. PrP aggregation lag time (n=6) dependence on GuHCl concentration (A). Optical density of S100A9 fibril samples (prepared in PBS without GuHCl, n=3) under a range of GuHCl concentrations (B). Aggregation kinetics of 50 μM and 200 μM S100A9 (n=4) under 0 M and 1.5 M GuHCl conditions (C). Fourier-transform infrared spectra of 200 μM S100A9 (non-aggregated control, yellow line) and its aggregates prepared under 0 M GuHCl conditions (control, black line), prepared under 0 M GuHCl and then incubated under 1.5 M GuHCl conditions (Inc, purple line) or prepared under 1.5 M GuHCl conditions (Prep, red line).. Atomic force microscopy images of 200 μM S100A9 sample aggregates prepared under 0 M GuHCl and incubated under 1.5 M conditions (E) or prepared under 1.5 M GuHCl conditions (F) and their cross-sectional height distribution (G, n=30). Error bars are for one standard deviation, box plots indicate the interquartile range.

Next, it was determined under which denaturant conditions S100A9 amyloid fibrils would remain stable for extended periods of time. For this reason, S100A9 aggregates (prepared in PBS at 37°C, 200 μM protein concentration) were combined with a range of different GuHCl concentration solutions and incubated at 37°C for 24 hours. The resulting solution optical densities (Figure 1B) were similar to the control samples up to 1.5 M, indicating the fibrils remain stable under these conditions. However, beyond 1.5 M, the optical density decreased and reached a plateau at ∼3.0 M GuHCl, suggesting dissociation of fibrils. Fitting the data with a Boltzmann sigmoidal equation revealed that the denaturation midpoint was at 2.2 M. Based on both of these assays, it was evident that a GuHCl concentration of 1.5 M would be a suitable intermediate point between a rapid onset of prion protein aggregation and the stability of S100A9 fibrils.

To examine how such a denaturant concentration would influence the aggregation process of native S100A9, its fibril formation kinetics were monitored under both 0 M and 1.5 M GuHCl conditions (Figure 1C). Without GuHCl, the aggregation of 200 μM S100A9 proceeded quickly with only a short lag phase and there was an overall 10-fold increase in thioflavin-T (ThT) fluorescence. In the presence of 1.5 M GuHCl, the fluorescence intensity value was significantly smaller, and the signal experienced a long, gradual increase over the entire monitoring period. Since ThT fluorescence intensity can be modulated by different solution ionic strength values, it can not be directly compared between conditions. For this reason, the concentration of aggregated S100A9 was determined as described in the Materials and Methods section. Both 0 M and 1.5 M GuHCl conditions resulted in residual, non-aggregated protein, however, its concentration was significantly higher in the case of 1.5 M GuHCl (∼65 μM, as opposed to ∼23 μM out of the initial 200 μM). At lower S100A9 concentration (50 μM) and under 1.5 M GuHCl conditions, there were no detectable signs of ThT fluorescence increase during this time period and the concentrations of non-aggregated S100A9 remained similar to its initial reaction solution (Figure 1C).These results displayed that such conditions (50 μM S100A9, pH 7.4, PBS with 1.5 M GuHCL) would be suitable to maintain S100A9 in its non-aggregated state for the duration of the experiments with PrP amyloid formation. The non-aggregated S100A9 solution was comprised of monomeric and dimeric forms of the protein (based on SDS-PAGE Supplementary Figure S2), however, it cannot be ruled out that higher oligomeric and ThT-negative species could also exist in the incubated solutions.

Prolonged incubation times under conditions with GuHCl may induce a certain level of structural or morphological alterations. To examine this possibility, S100A9 fibrils were initially prepared under 0 M GuHCl and then incubated under 1.5 M GuHCl conditions, or initially prepared under 1.5 M GuHCl conditions. They were then examined using Fourier-transform infrared spectroscopy (Figure 1D), as well as atomic force microscopy (AFM, Figure 1E, F). In the case of S100A9 fibrils prepared without GuHCl and then incubated with the denaturant, the FTIR spectrum retained a high level of similarity to the control (control did not change during incubation without GuHCl), with an identical main maximum position (1618 cm^−1^, related to strong beta-sheet hydrogen bonding ^55^) and a slightly lower ratio between the main maximum position and 1650 cm^−1^ (related to random-coil or alpha-helix hydrogen bonding). This suggests that, for the most part, S100A9 fibrils were capable of retaining their original secondary structure. For S100A9 aggregates which were prepared under 1.5 M GuHCl conditions, the main maximum was shifted towards 1619 cm^−1^, the 1618 cm^−1^ / 1650 cm^−1^ ratio was lower than both the control and the incubated fibril spectra ratios and the small peak associated with anti-parallel beta-sheets (1692 cm^−1^) was slightly shifted towards a higher wavenumber. Deconvolution of all three spectra confirmed that incubation of S100A9 fibrils at 1.5 M GuHCl had a lower effect on the reduction of beta-sheet content than when S100A9 aggregation was conducted under the same conditions (Supplementary Figure S3).

Analysis of both samples with AFM revealed that they also had distinct morphological characteristics. Both types of aggregates possessed the appearance of short, worm-like structures, which have been reported in previous studies ^56^. However, the fibrils prepared at 1.5 M GuHCl had a significantly higher average cross-sectional height distribution (Figure 1G, n=30, p<0.001).

Considering that the selected 1.5 M GuHCl conditions yielded relatively quick aggregation of PrP, maintained the stability of preformed S100A9 fibrils and inhibited the aggregation of native S100A9, they were selected for further cross-interaction experiments. Due to the statistically significant difference of S100A9 aggregates prepared under 1.5 M GuHCl conditions, they were also analysed alongside the 0 M GuHCl fibrils and non-aggregated S100A9.

To examine how the presence of S100A9 affects the aggregation of PrP, a range of non-aggregated and fibrillar S100A9 concentrations were combined with 25 μM prion protein and the fibril formation reaction was tracked in the presence of 1.5 M GuHCl, 600 RPM orbital agitation and a constant 37°C temperature. All statistical analysis was done using an ANOVA Bonferroni means comparison (n=16, p<0.001). When PrP was in the presence of non-aggregated S100A9, no statistically significant differences were observed at 1 μM, 2.5 μM and 5 μM protein concentrations, despite a slow and gradual increase in the process lag time values (Figure 2A, Supplementary Figure S4). Higher S100A9 concentrations, however, led to a significant increase in lag time when compared to the control and lower protein concentrations, with 25 μM conditions having an average value of 1140 minutes and 50 μM – 1760 minutes. Opposite to the effect of non-aggregated S100A9, the entire concentration range of its aggregated forms, prepared under 0 M GuHCl (Figure 2B) did not cause statistically significant deviations from the control sample lag time values. Similar results were also obtained when using S100A9 fibrils, which were prepared under 1.5 M GuHCl conditions (Figure 2C).

**Figure 2.**
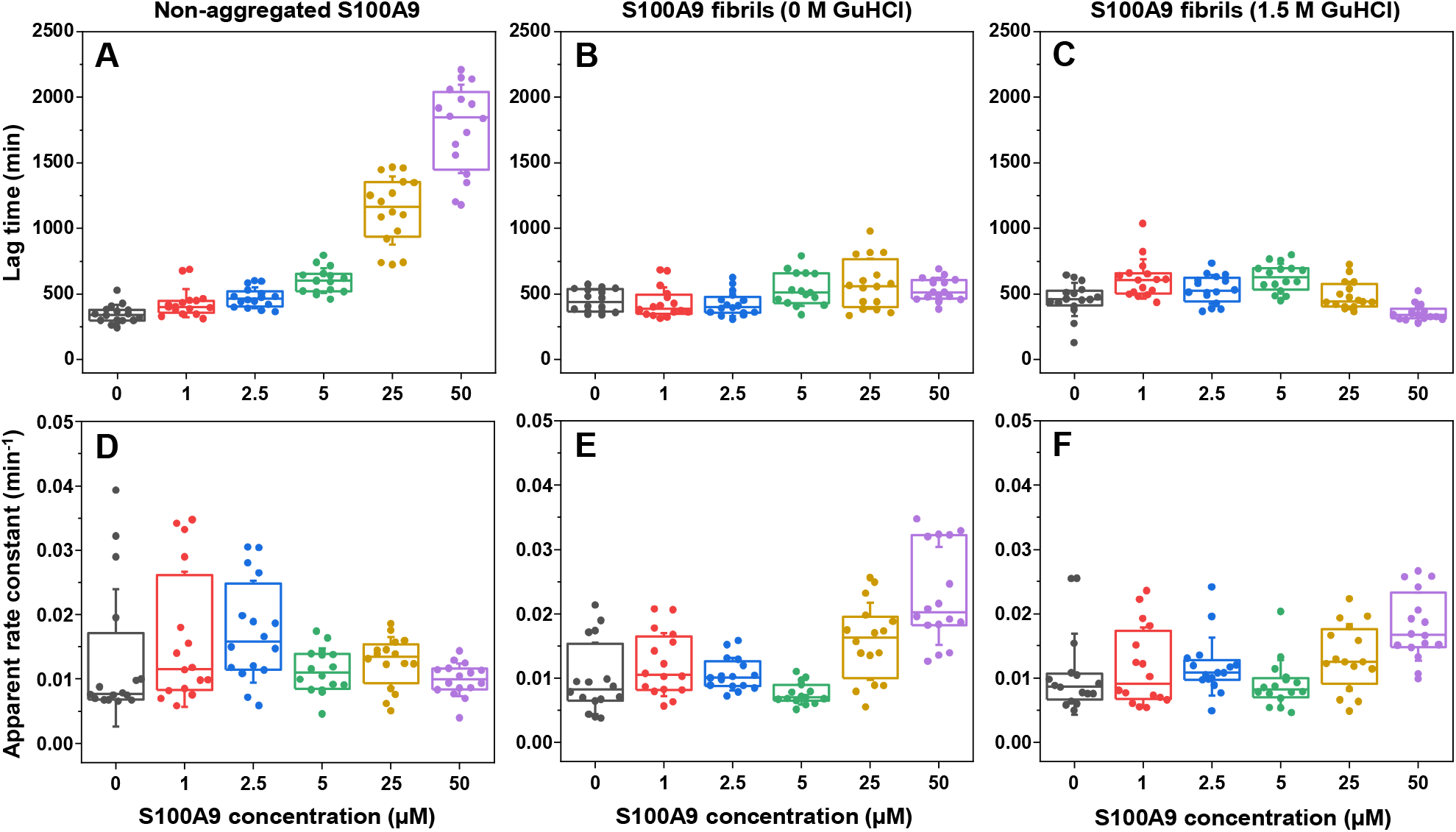
PrP aggregation kinetic parameters in the presence of different S100A9 concentrations. The lag time (A – C) and apparent rate constant (D – F) distributions of prion protein aggregation in the presence of non-aggregated S100A9 (first column), S100A9 fibrils prepared under 0 M GuHCl (second column) and 1.5 M conditions (third column). All data points are colour-coded, error bars are for one standard deviation (n=16) and box plots indicate the interquartile range.

Contrary to the influence that non-aggregated S100A9 had on the lag time of PrP aggregation, it did not cause significant changes to the reaction’s apparent rate constant (Figure 2D). However, there was a notable reduction in the process stochasticity, with 5, 25 and 50 μM concentrations leading to a reduced variation in the apparent rate constant. In the case of S100A9 fibrils prepared under 0 M GuHCl conditions, only the highest S100A9 concentration yielded apparent rate values that were significantly higher than the control (Figure 2E). A peculiar aspect was observed for the 5 μM S100A9 concentration, where it pertained a relatively low level of stochasticity when using all three forms of S100A9. The 1.5 M GuHCl S100A9 aggregates did not have such a profound effect, which may be related to either their different structure or the presence of a substantial concentration of non-aggregated S100A9 (Figure 2F).

Finally, it was investigated how the presence of S100A9 modulated PrP aggregate ThT-binding properties and fluorescence of the samples, as these parameters may be indicative of different PrP fibril strain formation ^57^. When using non-aggregated S100A9, only the 50 μM concentration conditions yielded a significantly lower average fluorescence signal intensity, while all other conditions remained similar to the control (Supplementary Figure S5A). S100A9, which was aggregated under 0 M GuHCl conditions, on the other hand, had significantly diminished fluorescence intensities under 1, 2.5 and 5 μM S100A9 concentrations (Supplementary Figure S5B). Interestingly, the presence of 25 and 50 μM S100A9 resulted in aggregates with similar values as the control, albeit with a massively reduced stochasticity. As was the case with the apparent rate constant results, the 1.5 M GuHCl S100A9 fibrils led to an intermediate variant of intensity values, with part of them being similar to the non-aggregated S100A9 and others – to the 0 M S100A9 fibril conditions (Supplementary Figure S5C). Unlike the 0 M GuHCl aggregates, the high level of stochasticity for most of the conditions yielded no statistically significant deviations from the control. One notable observation was the highly reduced stochasticity of the 50 μM S100A9 sample fluorescence intensities when using all three forms of S100A9, similar to the case of the apparent rate constant at 5 μM protein concentration.

S100A9 is a calcium-binding protein, which is a factor that may influence its interaction with PrP. In order to examine this possibility, the kinetics of PrP amyloid formation were monitored with an additional 0.5 mM CaCl_2_ in the reaction solutions (10-times the concentration of S100A9). A comparison of the control and conditions with 50 μM non-aggregated S100A9 revealed that the effect of S100A9 remained similar as in the absence of calcium ions (Supplementary Figure 6). The only notable distinction was the slightly higher lag phase for both conditions, likely as a result of the additional CaCl_2_. These results indicate that either calcium binding does not play a role in modulating the aggregation of PrP or that the presence of a denaturant destabilizes the S100A9-calcium interactions.

To further explore the influence of S100A9 on the secondary structure of PrP aggregates and whether they are capable of self-replication the samples were reseeded and then analysed using FTIR spectroscopy. Since most of them contained mixtures of PrP and S100A9, the samples were subjected to two rounds of reseeding, where aliquots of each sample (40 μL) were combined with the initial aggregation solution (360 μL, which did not contain S100A9) and incubated under identical conditions as were used in the initial kinetic monitoring assay. This procedure was done to reduce the concentration of S100A9 in the sample to negligible levels, determine if the PrP aggregates are capable of self-replication, and to reduce the possible polymorphism of PrP fibrils ^58^It was observed that the seeded aggregation reactions proceeded without a lag-phase and reached a plateau in less than 10 hours, indicating effective self-replication (Supplementary Figures S7 and S8). Taking into consideration that both the 0 M and 1.5 M GuHCl condition S100A9 fibrils previously displayed a similar influence on PrP aggregation kinetic parameters, only the 0 M condition samples were analysed in the FTIR assay.

Since even after two rounds of reseeding the prion protein aggregates may possess slightly varying secondary structures ^57^, all 16 FTIR spectra and their second derivatives from each condition were superimposed and the dominant types were used for comparison. The FTIR spectra of the control samples (Figure 3A) had main maxima positions at 1627 – 1628 cm^−1^ and their second derivatives displayed minima at 1611 cm^−1^ and 1627 cm^−1^, suggesting two types of hydrogen bonding in the beta-sheet structure. The second derivatives also contained minima at 1647 cm^−1^ (alpha-helix or random coils), as well as 1659, 1668 and 1677 cm^−1^ (turn/loop motifs). Spectra of samples with an initial 5 μM non-aggregated S100A9 were identical (Figure 3B), with similar minima positions as the control samples (Figure 3 table insert). In the case of 5 μM S100A9 fibrils, however, the dominant PrP FTIR spectra second derivatives (Figure 3C) displayed two minima at 1616 and 1628 cm^−1^, suggesting the formation of different strength beta-sheet hydrogen bonding.

**Figure 3.**
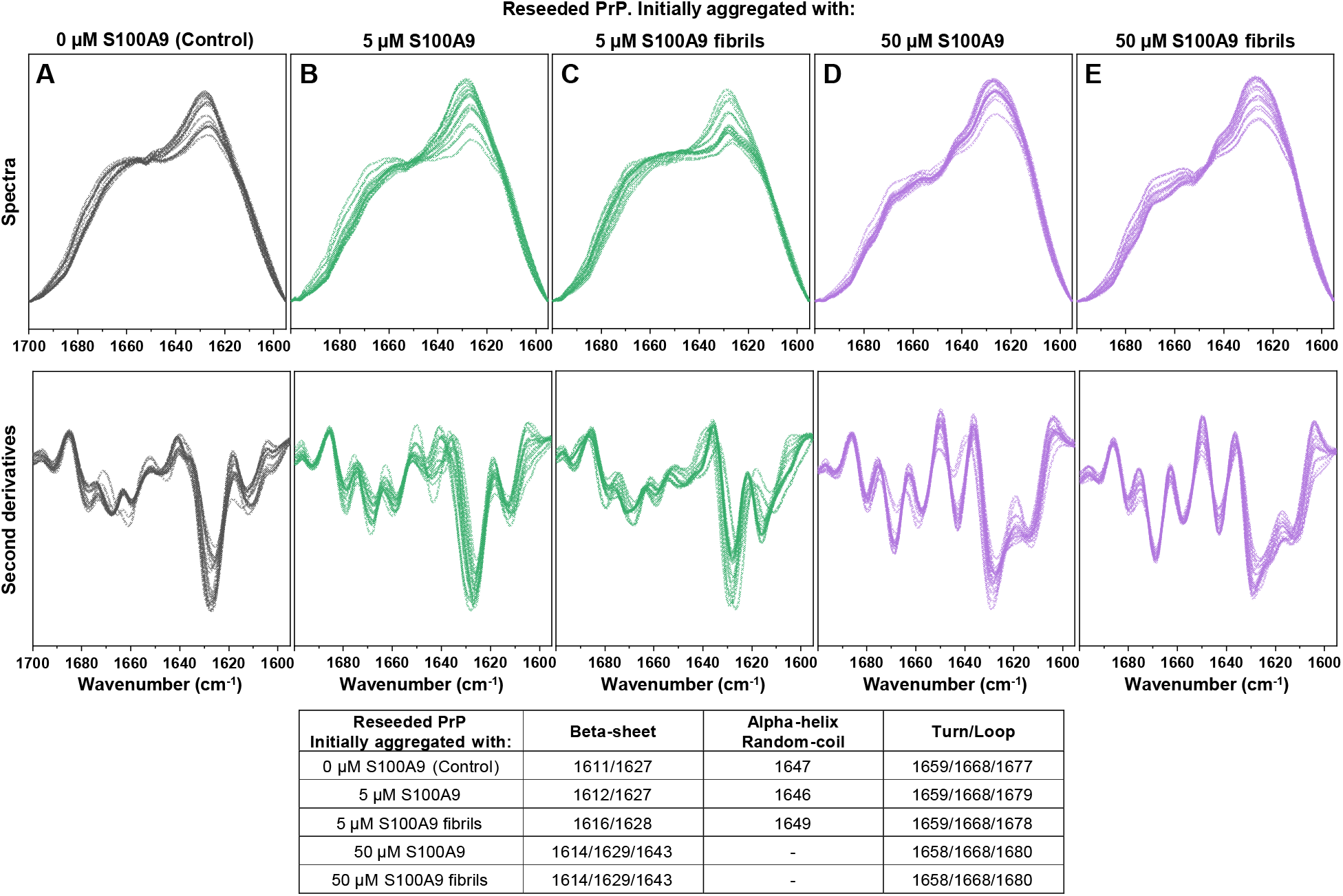
Secondary structure comparison of reseeded PrP fibrils formed in the presence of different S100A9 concentrations. The FTIR spectra of reseeded PrP fibrils, initially formed in the absence (A, control) and in the presence of 5 μM non-aggregated S100A9 (B), 5 μM S100A9 fibrils (C), 50 μM non-aggregated S100A9 (D) and 50 μM S100A9 fibrils (E)and their second derivatives. Due to two rounds of reseeding and 100-time S100A9 dilution, the spectra are representative of only the PrP structure. Table insert displays the dominant spectra second derivative minima positions related to beta-sheet, alpha-helix, random-coil and turn/loop hydrogen bonding.

When 50 μM of S100A9 was present during the aggregation of PrP, both their forms induced the formation of a different type of PrP amyloid structure. The resulting FTIR spectra (Figure 3D, E) main maxima were shifted to 1626 cm^−1^, while the second derivatives contained minima at 1614, 1629 and 1643 cm^−1^ (associated with three different strength hydrogen bonds in the beta-sheet structure). The second derivatives also did not contain a minimum at ∼1650, implying a lower amount of alpha-helical or random-coil regions. The positions associated with turn/loop motifs remained fairly similar to the control and 5 μM S100A9 condition spectra. Interestingly, the various spectra observed during this study were almost identical to ones determined in a previous amyloid protein interaction study involving PrP and superoxide dismutase-1 (SOD1). There, the presence of 25 μM SOD1 resulted in PrP aggregates with FTIR spectra similar to ones formed under 50 μM S100A9 conditions^46^. The previous study also revealed that both the control, as well as 5 μM and 50 μM S100A9 condition PrP structures had a high level of self-association, resulting in the assembly of large aggregate clumps. Analysis of their morphology also indicated that the 50 μM S100A9 condition PrP aggregate type had a slightly higher cross-sectional height^46^.

Since the presence of S100A9 had a significant effect on PrP spontaneous aggregation kinetics and the resulting fibril secondary structure, it was further investigated whether S100A9 also affected their self-replication properties. Different concentrations of non-aggregated and fibrillar S100A9 were incubated with the initial prion protein reaction solutions, containing 10% of PrP fibrils. In the absence of S100A9, the reaction proceeded without a lag phase and the average aggregation half-time value (t_50_) was 160 minutes (Figure 4). The presence of S100A9 resulted in sigmoidal aggregation curves with higher t_50_ values, suggesting that it may be capable of inhibiting the self-replication properties of PrP fibrils. In the case of non-aggregated S100A9 (Figure 4A), there was a concentration-dependent increase in t_50_ values up to 40 μM (Figure 4C). Oppositely, the inhibitory effect became saturated at only 20 μM of S100A9 fibrils (Figure 4B, C). A comparison of PrP aggregation half-time values in the presence of 50 μM S100A9 (Figure 4C) revealed that the inhibition was higher when S100A9 was in its non-aggregated state.

**Figure 4.**
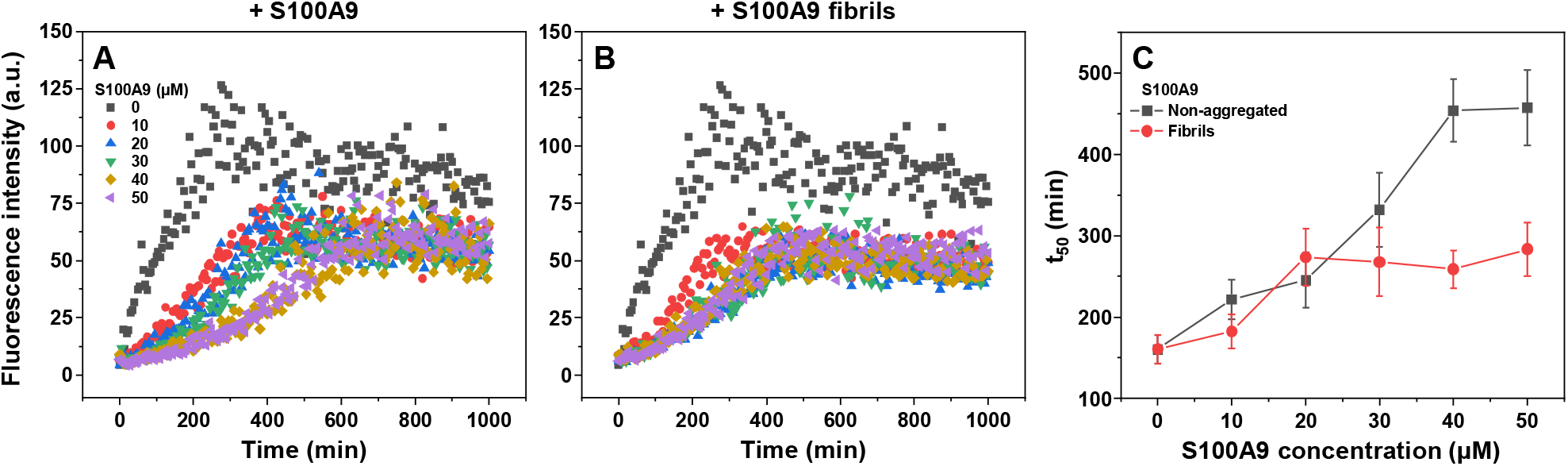
Seeded aggregation of prion proteins in the presence of S100A9. Examples of PrP aggregation curves in the presence of 10% PrP seed with different concentrations of non-aggregated S100A9 (A) and S100A9 fibrils (B). Aggregation curve half-time values (t_50_, C). Error bars are for one standard deviation (n=6).

The resulting aggregate samples were then examined by FTIR spectroscopy without any additional reseeding steps. Since 50 μM S100A9 under 1.5 M GuHCl conditions could not form detectable amounts of aggregates (Figure 1C), the spectra of samples which contained non-aggregated S100A9 could be compared without any need to account for the protein’s presence (S100A9 was removed during the H_2_O/D_2_O exchange). In this case, the spectra main maximum was similar for all the samples (1627 – 1628 cm^−1^), while there was a notable difference between the peak sizes associated with beta-sheet hydrogen bonding. There appeared to be a S100A9-concentration-dependent increase in the relative height of the band at 1627-1628 cm^−1^, when compared to the random-coil, alpha-helix and turn/loop regions (1650 – 1680 cm^−1^). These results suggested that S100A9 may have inhibited the self-replication of certain fibril types or promoted the formation of aggregates with larger beta-sheet content in their structure.

FTIR analysis of samples where S100A9 was in its fibrillar state was more complicated, as both protein aggregates were present in the sample. Surprisingly, it was discovered that the spectra from both sets of conditions can be deconvoluted into spectra of fibrillar S100A9, PrP aggregates formed without S100A9 (containing a relatively lower content of beta-sheets, referred to as type I, Figure 5A bottom) and PrP aggregates formed in the presence of 50 μM non-aggregated S100A9 (containing a relatively higher content of beta-sheets, referred to as type II, Figure 5A top). This deconvolution procedure (Supplementary Figure S9) revealed that higher concentrations of both types of S100A9 promoted the formation of type II PrP fibrils. In both cases, the 50 μM S100A9 conditions yielded identical PrP fibril secondary structures, regardless whether S100A9 was in its non-aggregated or fibrillar state.

**Figure 5.**
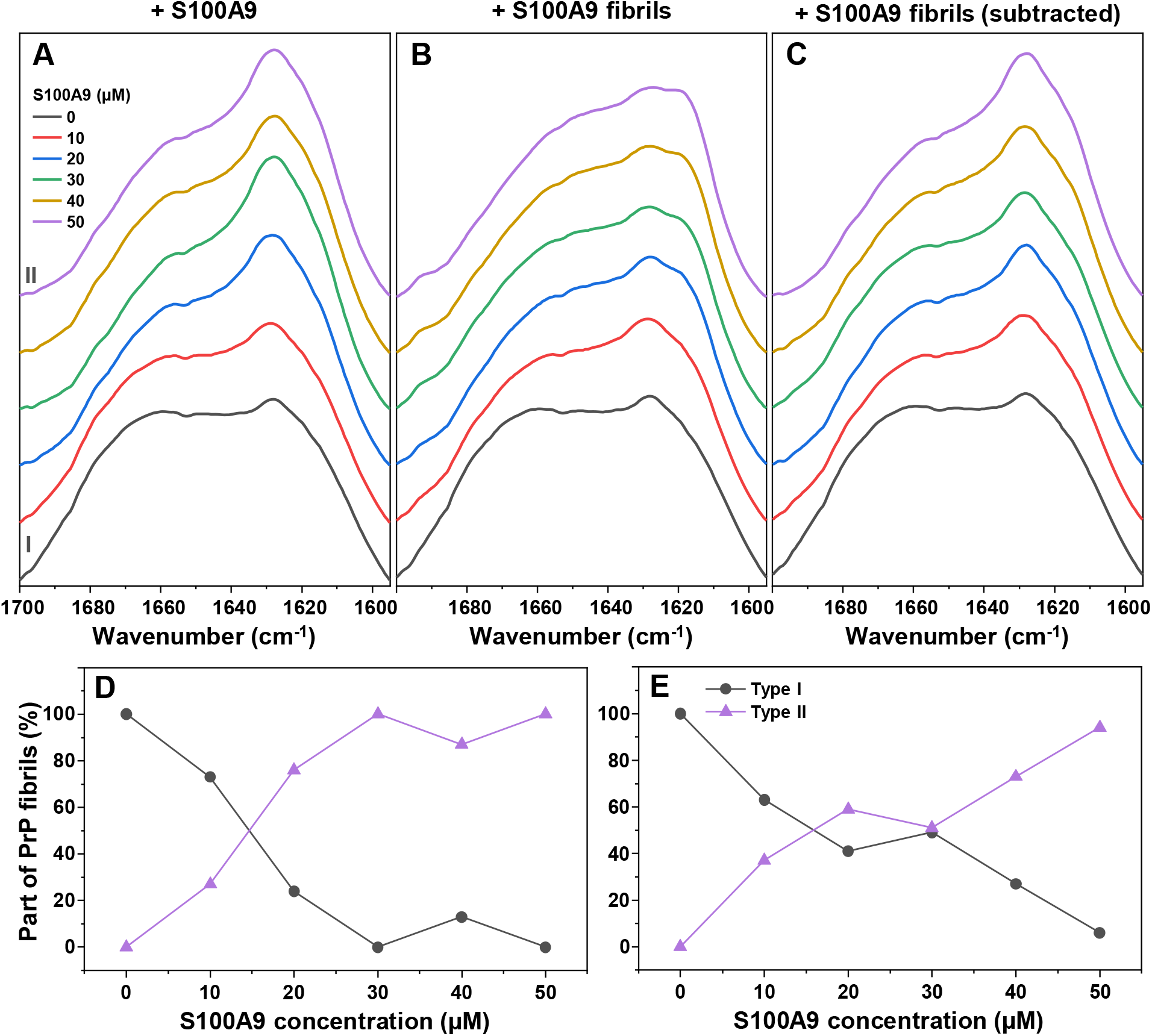
Influence of S100A9 on PrP fibril secondary structures during self-replication. Fourier-transform infrared spectra of PrP fibrils after self-replication in the presence of non-aggregated S100A9 (A) and S100A9 fibrils (B). Roman numerals mark type I and type II PrP fibrils. Panel B spectra after S100A9 spectra subtraction (C). FTIR spectra deconvolution results when PrP was replicated in the presence of non-aggregated S100A9 (D) or S100A9 fibrils (E). Deconvolution of each spectrum into the aggregated S100A9 and PrP fibril types I and II is shown as Supplementary Figure S9. Deconvolution results presented in Panels D and E were calculated from single bulk solution FTIR spectra of 6 combined samples for each condition.

Based on the PrP fibril FTIR spectra in the presence of non-aggregated S100A9 (Figure 5A), there were no detectable signs of association between the two proteins (no appearance of peaks associated with non-aggregated S100A9), suggesting that the interaction may occur with monomeric PrP. During a protein melt assay under 0 M GuHCl conditions, it was observed that both proteins had very similar melting point temperature (T_m_) values, which made it difficult to distinguish the effect of their cross-interaction (Figure 6A). However, the melt curve when both proteins were in solution had an almost identical first derivative maximum position to the combined PrP and S100A9 melt curve (Figure 6A, inset). This result indicated that the presence of S100A9 did not alter the stability of monomeric PrP. Examination of the samples using size-exclusion chromatography also revealed that there were no notable interactions between the two proteins. The monomeric PrP elution peak (Figure 6B, C) was slightly shifted compared to the combined S100A9-PrP sample peak (10.50 mL and 10.37 mL respectively), indicating that the proteins did not form a complex in solution, or the interaction was transient.

**Figure 6.**
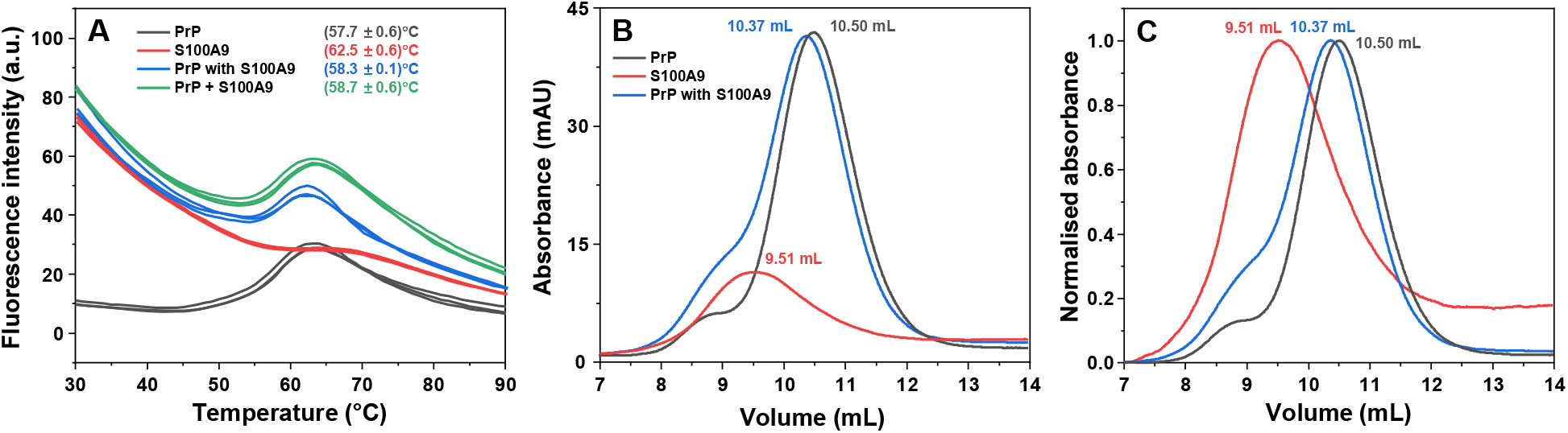
Examination of non-aggregated prion protein (PrP) and S100A9 cross-interactions. Protein melt assay of PrP and S100A9 (A, black line – PrP alone, red line – S100A9 alone, blue line – PrP and S100A9 together in solution, green line – combined PrP and S100A9 fluorescence intensities). Fluorescence curve first derivative maximum positions are color-coded and displayed in panel A. Size-exclusion chromatography elution profiles of PrP (black lines), S100A9 (red lines) and when they are both in solution (blue lines) (B – raw data, C - normalised).

Finally, it was investigated whether the different secondary structure PrP aggregates possessed distinct toxicities against neuroblastoma cells. Aggregates from replicated control PrP samples, as well as from samples initially prepared with 50 μM non-aggregated S100A9 were resuspended into PBS and applied to SH-SY5Y cells. MTT cell viability and LDH release assays (Supplementary Figure S10) revealed that there were no significant differences between the two different secondary structure PrP aggregates. These results indicated that the redirected pathway of PrP amyloid formation did not yield a less cytotoxic variant of aggregates.

## Discussion

In recent years, there has been a surge in the number of reports describing the cross-interaction of amyloidogenic proteins and how it influences their aggregation process. Both S100A9, as well as prion proteins have previously been shown to participate in these type of interactions ^22,29^, however, there was lack of information on how they may influence each other’s amyloid formation. Since S100A9 is a pro-inflammatory protein, which shares a localisation with PrP ^40–42^, this work was dedicated to gaining a better understanding of how it can alter the pathway of PrP aggregation *in vitro*.

Based on the non-seeded kinetics of PrP amyloid formation, the non-aggregated S100A9 had a significant inhibitory effect on the primary nucleation phase, while its fibrillar form did not alter the length of the lag phase. This suggests that non-aggregated S100A9 may be capable of disrupting the process of PrP amyloid nuclei formation, either by interacting with monomeric PrP or the prefibrillar structure. Analysis of S100A9 and PrP interactions by size-exclusion chromatography and a protein melt assay did not show any significant signs of complex formation when both proteins were in their native state and their association may have only been transient. The FTIR spectra of PrP structures formed in the presence of non-aggregated S100A9 also displayed a lack of cross-interaction between the two proteins when PrP was in its aggregated state. These observations further support the idea that S100A9 only interacts with the prefibrillar nuclei of PrP.

Oppositely to the effect on PrP aggregation lag time, the fibrillar form of S100A9 had a significant effect on the process apparent rate constant, resulting in a 2-fold increase. This result can be explained by the hydrophobic fibrillar surface of S100A9 acting as a catalyst for surface-mediated nucleation ^59^. Such an event may also contribute to the reduction of primary nucleation lag time and compete with the inhibitory effect of the non-aggregated form of S100A9. Since it is known that S100A9 is capable of spontaneously undergoing aggregation under physiological conditions, it is possible that this transition is at least partially responsible for the onset of prionopathies. Both proteins share a localization in the cerebral cortex ^40,41^ and cerebrospinal fluid ^42^, where native S100A9 would be able to prevent the nucleation process of PrP. As it forms amyloid-like fibrils itself, the inhibitory effect on PrP nuclei formation would cease and the aggregates would also serve as a surface for additional nucleation effects.

The complexity of the possible cross-interaction increased even further after an examination of how S100A9 affected the self-replication process of PrP fibrils. When the initial PrP aggregates were prepared in the absence of S100A9, they were also only effective at replicating their structure when there was no S100A9 present in the reaction solution. In this case, both the non-aggregated and fibrillar forms of S100A9 significantly increased the process half-time values and changed the PrP aggregation curve shapes from exponential to sigmoidal. The resulting PrP fibril secondary structure also became different than the initial “seed”, suggesting that the presence of S100A9 may facilitate the propagation of only certain PrP aggregate types, while inhibiting others. A similar effect was previously observed with alpha-synuclein, where the presence of S100A9 modulated the aggregation reaction and promoted the formation of a single alpha-synuclein fibril conformation ^47^.

Further support for the idea of S100A9 selectively altering the amyloid formation of PrP could be obtained by analysing the PrP fibril structures generated during spontaneous aggregation. When the reaction solutions contained a small concentration of S100A9, some of the samples resulted in FTIR spectra similar to the ones observed in the self-replication experiment. At higher S100A9 concentrations, the PrP fibril FTIR spectra became completely different from the controls, while being nearly identical between non-aggregated and fibrillar S100A9 conditions. In our previous examination of PrP and superoxide dismutase-1 (SOD1) cross-interaction, similar FTIR spectra were also observed when PrP was aggregated in the presence of SOD1 ^46^. Under 50 μM SOD1 conditions, however, the FTIR spectra were completely different from the ones observed under 50 μM S100A9 conditions, indicating that different amyloidogenic proteins may have distinct effects on PrP aggregation.

Another peculiar aspect was the difference between the secondary structure of PrP aggregates depending on whether they initially formed in the presence of S100A9 or were only replicated with S100A9. This observation implies that the cross-interaction promotes the formation of certain PrP amyloid nuclei, which can effectively replicate under conditions with S100A9. However, if the reaction solution is “seeded” with preformed PrP aggregates, S100A9 inhibits their elongation or only facilitates the replication of specific fibril types. These results further support the idea that S100A9 interacts with the aggregation centers of PrP, whether they are the prefibrillar nuclei or the elongation-competent ends of fibrils.

The results of this study demonstrate the highly complex cross-interaction between the proinflammatory S100A9 and prion proteins during amyloid aggregation. S100A9 is capable of altering both the kinetic parameters of PrP fibril formation, as well as their resulting secondary structure and self-replication properties. Taking into consideration the shared localisation with PrP and previous reports of possible S100A9 involvement in neurodegenerative disorders, this study adds another piece to the complex amyloid interactome puzzle.

## Conclusions

The cross-interaction between S100A9 and PrP leads to a redirection of the PrP aggregation pathway and a significant inhibition of the primary nucleation phase. When S100A9 is in its fibrillar form, however, the inhibitory effect is diminished and the aggregate surface acts as a catalyst for PrP secondary nucleation. These results, along with previously observed S100A9 cross-interactions, demonstrate its potential role against the onset of neurodegenerative disorders.

## Author Contributions

Conceptualization, M.Z. and V.S.; investigation, M.Z., D.S., K.M., A.S. and R.S.; resources, V.S.; formal analysis, M.Z., D.S. and V.S.; writing—original draft preparation, M.Z.; writing— review and editing, M.Z., D.S., K.M., A.S. and V.S. All authors have read and agreed to the published version of the manuscript.

## Data Availability Statement

All data are available in the Mendeley data online repository: https://data.mendeley.com/datasets/8t7hxxnc53/1

## References

1. Chiti, F. & Dobson, C. M. Protein Misfolding, Amyloid Formation, and Human Disease: A Summary of Progress Over the Last Decade. Annu. Rev. Biochem. 86, 27–68 (2017).

2. Knowles, T. P. J., Vendruscolo, M. & Dobson, C. M. The amyloid state and its association with protein misfolding diseases. Nat. Rev. Mol. Cell Biol. 15, 384–396 (2014).

3. Iadanza, M. G., Jackson, M. P., Hewitt, E. W., Ranson, N. A. & Radford, S. E. A new era for understanding amyloid structures and disease. Nat. Rev. Mol. Cell Biol. 19, 755–773 (2018).

4. Lövestam, S. et al. Assembly of recombinant tau into filaments identical to those of Alzheimer’s disease and chronic traumatic encephalopathy. Elife 11, 1–27 (2022).

5. Yang, Y. et al. Structures of α-synuclein filaments from human brains with Lewy pathology. Nature 610, 791–795 (2022).

6. Cummings, J. et al. Alzheimer’s disease drug development pipeline: 2023. Alzheimer’s Dement. Transl. Res. Clin. Interv. 9, (2023).

7. Mehta, D., Jackson, R., Paul, G., Shi, J. & Sabbagh, M. Why do trials for Alzheimer’s disease drugs keep failing? A discontinued drug perspective for 2010-2015. Expert Opin. Investig. Drugs 26, 735–739 (2017).

8. Arthur, K. C. et al. Projected increase in amyotrophic lateral sclerosis from 2015 to 2040. Nat. Commun. 7, 12408 (2016).

9. Brookmeyer, R., Gray, S. & Kawas, C. Projections of Alzheimer’s disease in the United States and the public health impact of delaying disease onset. Am. J. Public Health 88, 1337–1342 (1998).

10. Meisl, G. et al. Molecular mechanisms of protein aggregation from global fitting of kinetic models. Nat. Protoc. 11, 252–272 (2016).

11. Meisl, G. et al. Differences in nucleation behavior underlie the contrasting aggregation kinetics of the Aβ40 and Aβ42 peptides. Proc. Natl. Acad. Sci. 111, 9384–9389 (2014).

12. Buell, A. K. et al. Solution conditions determine the relative importance of nucleation and growth processes in-synuclein aggregation. Proc. Natl. Acad. Sci. 111, 7671–7676 (2014).

13. Gurry, T. & Stultz, C. M. Mechanism of amyloid-β fibril elongation. Biochemistry 53, 6981–6991 (2014).

14. Rodriguez, R. A., Chen, L. Y., Plascencia-Villa, G. & Perry, G. Thermodynamics of Amyloid-β Fibril Elongation: Atomistic Details of the Transition State. ACS Chem. Neurosci. 9, 783–789 (2018).

15. Nicoud, L., Lazzari, S., Balderas Barragán, D. & Morbidelli, M. Fragmentation of Amyloid Fibrils Occurs in Preferential Positions Depending on the Environmental Conditions. J. Phys. Chem. B 119, 4644–4652 (2015).

16. Törnquist, M. et al. Secondary nucleation in amyloid formation. Chem. Commun. 54, 8667–8684 (2018).

17. Gaspar, R. et al. Secondary nucleation of monomers on fibril surface dominates α-synuclein aggregation and provides autocatalytic amyloid amplification. Q. Rev. Biophys. 50, (2017).

18. Wiśniewski, J. R., Hein, M. Y., Cox, J. & Mann, M. A ‘proteomic ruler’ for protein copy number and concentration estimation without spike-in standards. Mol. Cell. Proteomics 13, 3497–3506 (2014).

19. Ono, K., Takahashi, R., Ikeda, T. & Yamada, M. Cross-seeding effects of amyloid β-protein and α-synuclein. J. Neurochem. 122, 883–890 (2012).

20. Vasconcelos, B. et al. Heterotypic seeding of Tau fibrillization by pre-aggregated Abeta provides potent seeds for prion-like seeding and propagation of Tau-pathology in vivo. Acta Neuropathol. 131, 549–569 (2016).

21. Nisbet, R. M., Polanco, J.-C., Ittner, L. M. & Götz, J. Tau aggregation and its interplay with amyloid-β. Acta Neuropathol. 129, 207–220 (2015).

22. Honda, R. Amyloid-β Peptide Induces Prion Protein Amyloid Formation: Evidence for Its Widespread Amyloidogenic Effect. Angew. Chemie - Int. Ed. 57, 6086–6089 (2018).

23. Guo, J. L. et al. Distinct α-Synuclein Strains Differentially Promote Tau Inclusions in Neurons. Cell 154, 103–117 (2013).

24. Lu, J. et al. Structural basis of the interplay between α-synuclein and Tau in regulating pathological amyloid aggregation. J. Biol. Chem. 295, 7470–7480 (2020).

25. Katorcha, E. et al. Cross-seeding of prions by aggregated α-synuclein leads to transmissible spongiform encephalopathy. PLoS Pathog. 13, 1–23 (2017).

26. Burdukiewicz, M. et al. AmyloGraph: a comprehensive database of amyloid–amyloid interactions. Nucleic Acids Res. 51, D352–D357 (2023).

27. Vogl, T., Gharibyan, A. L. & Morozova-Roche, L. A. Pro-inflammatory S100A8 and S100A9 proteins: Self-assembly into multifunctional native and amyloid complexes. Int. J. Mol. Sci. 13, 2893–2917 (2012).

28. Wang, C. et al. The role of pro-inflammatory S100A9 in Alzheimer’s disease amyloid-neuroinflammatory cascade. Acta Neuropathol. 127, 507–522 (2014).

29. Horvath, I. et al. Co-aggregation of pro-inflammatory S100A9 with α-synuclein in Parkinson’s disease: ex vivo and in vitro studies. J. Neuroinflammation 15, 172 (2018).

30. Esposito, G. et al. S100B induces tau protein hyperphosphorylation via Dickopff-1 up-regulation and disrupts the Wnt pathway in human neural stem cells. J. Cell. Mol. Med. 12, 914–927 (2008).

31. Iashchishyn, I. A., Sulskis, D., Nguyen Ngoc, M., Smirnovas, V. & Morozova-Roche, L. A. Finke– Watzky Two-Step Nucleation–Autocatalysis Model of S100A9 Amyloid Formation: Protein Misfolding as “Nucleation” Event. ACS Chem. Neurosci. 8, 2152–2158 (2017).

32. Linden, R. The biological function of the prion protein: A cell surface scaffold of signaling modules. Front. Mol. Neurosci. 10, 1–19 (2017).

33. Collinge, J. Variant Creutzfeldt-Jakob disease. Lancet 354, 317–323 (1999).

34. Baldwin, K. J. & Correll, C. M. Prion Disease. Semin. Neurol. 39, 428–439 (2019).

35. Caughey, B. & Chesebro, B. Transmissible spongiform encephalopathies and prion protein interconversions. Adv. Virus Res. 56, (2001).

36. Bousset, L. et al. Structural and functional characterization of two alpha-synuclein strains. Nat. Commun. 4, 2575 (2013).

37. Yang, S. et al. Polymorphisms at amino acid residues 141 and 154 influence conformational variation in ovine PrP. Biomed Res. Int. 2014, 372491 (2014).

38. Fändrich, M., Meinhardt, J. & Grigorieff, N. Structural polymorphism of Alzheimer Aβ and other amyloid fibrils. Prion 3, 89–93 (2009).

39. Wang, M., Herrmann, C. J., Simonovic, M., Szklarczyk, D. & von Mering, C. Version 4.0 of PaxDb: Protein abundance data, integrated across model organisms, tissues, and cell-lines. Proteomics 15, 3163–3168 (2015).

40. Kim, M.-S. et al. A draft map of the human proteome. Nature 509, 575–581 (2014).

41. Bauernfeind, A. L. et al. High spatial resolution proteomic comparison of the brain in humans and chimpanzees. J. Comp. Neurol. 523, 2043–2061 (2015).

42. Guldbrandsen, A. et al. In-depth Characterization of the Cerebrospinal Fluid (CSF) Proteome Displayed Through the CSF Proteome Resource (CSF-PR). Mol. Cell. Proteomics 13, 3152–3163 (2014).

43. Li, B., Chen, M. & Zhu, C. Neuroinflammation in Prion Disease. Int. J. Mol. Sci. 22, 2196 (2021).

44. Sanders, E. et al. The Stabilization of S100A9 Structure by Calcium Inhibits the Formation of Amyloid Fibrils. Int. J. Mol. Sci. 24, (2023).

45. Sorrentino, S. et al. Calcium Binding Promotes Prion Protein Fragment 90–231 Conformational Change toward a Membrane Destabilizing and Cytotoxic Structure. PLoS One 7, e38314 (2012).

46. Ziaunys, M., Mikalauskaite, K., Veiveris, D., Sakalauskas, A. & Smirnovas, V. Superoxide dismutase-1 alters the rate of prion protein aggregation and resulting fibril conformation. Arch. Biochem. Biophys. 715, 109096 (2022).

47. Toleikis, Z. et al. S100A9 Alters the Pathway of Alpha-Synuclein Amyloid Aggregation. Int. J. Mol. Sci. 22, 7972 (2021).

48. Smirnovas, V. et al. Structural organizaton of brain-derived mammalian prions as probed by hydrogen excange. Nat Struct Mol Biol 18, 504–506 (2011).

49. Milto, K., Michailova, K. & Smirnovas, V. Elongation of Mouse Prion Protein Amyloid-Like Fibrils: Effect of Temperature and Denaturant Concentration. PLoS One 9, e94469 (2014).

50. Mikalauskaite, K., Ziaunys, M., Sneideris, T. & Smirnovas, V. Effect of Ionic Strength on Thioflavin-T Affinity to Amyloid Fibrils and Its Fluorescence Intensity. Int. J. Mol. Sci. 21, 8916 (2020).

51. Foderà, V. et al. Thioflavin T Hydroxylation at Basic pH and Its Effect on Amyloid Fibril Detection. J. Phys. Chem. B 112, 15174–15181 (2008).

52. Mikalauskaite, K., Ziaunys, M. & Smirnovas, V. Lysozyme Amyloid Fibril Structural Variability Dependence on Initial Protein Folding State. Int. J. Mol. Sci. 23, 5421 (2022).

53. Nečas, D. & Klapetek, P. Gwyddion: An open-source software for SPM data analysis. Cent. Eur. J. Phys. 10, 181–188 (2012).

54. Sakalauskas, A. et al. Exploring the Formation of Polymers with Anti-Amyloid Properties within the 2131-Dihydroxyflavone Autoxidation Process. Antioxidants 11, (2022).

55. Barth, A. Infrared spectroscopy of proteins. Biochim. Biophys. Acta - Bioenerg. 1767, 1073–1101 (2007).

56. Pansieri, J. et al. Templating S100A9 amyloids on Aβ fibrillar surfaces revealed by charge detection mass spectrometry, microscopy, kinetic and microfluidic analyses. Chem. Sci. 11, 7031–7039 (2020).

57. Ziaunys, M., Sneideris, T. & Smirnovas, V. Formation of distinct prion protein amyloid fibrils under identical experimental conditions. Sci. Rep. 10, 4572 (2020).

58. Sun, Y. et al. Direct Observation of Competing Prion Protein Fibril Populations with Distinct Structures and Kinetics. ACS Nano 17, 6575–6588 (2023).

59. Koloteva-Levine, N. et al. Amyloid particles facilitate surface-catalyzed cross-seeding by acting as promiscuous nanoparticles. Proc. Natl. Acad. Sci. U. S. A. 118, 1–12 (2021).

