## Supplementary Material for "S100A9 Inhibits and Redirects Prion Protein 89-230 Fragment Amyloid Aggregation"

### Supplementary information

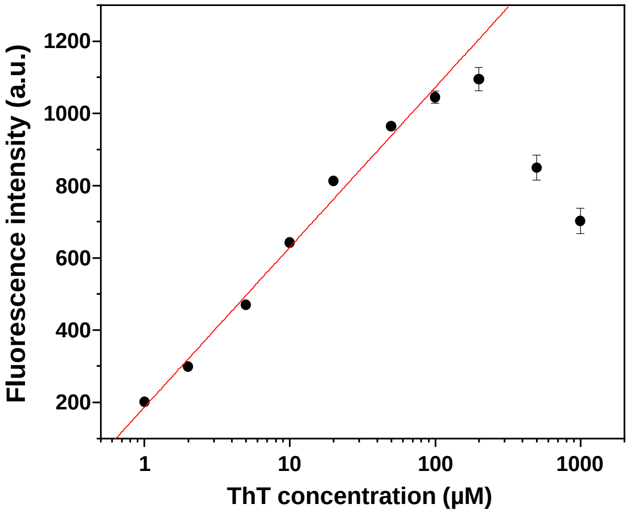


**Figure S1.** Thioflavin-T (ThT) fluorescence intensity dependence on ThT concentration. Dye fluorescence was measured using a ClarioStar Plus platereader, 10 minutes after combining prion protein (PrP) fibril samples with different concentrations of ThT at 22°C (average of 3 technical replicates for each condition, error bars are for one standard deviation). Final solutions contained 1.5 M GuHCl, 1 – 1000 µM ThT and 20 µM PrP fibrils in PBS (pH 7.4).

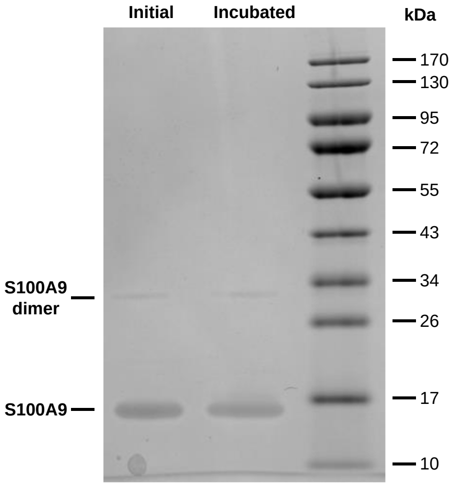


**Figure S2.** SDS-PAGE gel of 50 µM S100A9 before and after incubation in PBS containing 1.5 M GuHCl at 37°C. Before loading to the gel, soluble protein samples were mixed with Laemmli sample buffer and denatured by heating. EZ-Run™ Prestained *Rec* protein ladder (Fisher Scientific) was used as molecular weight markers.

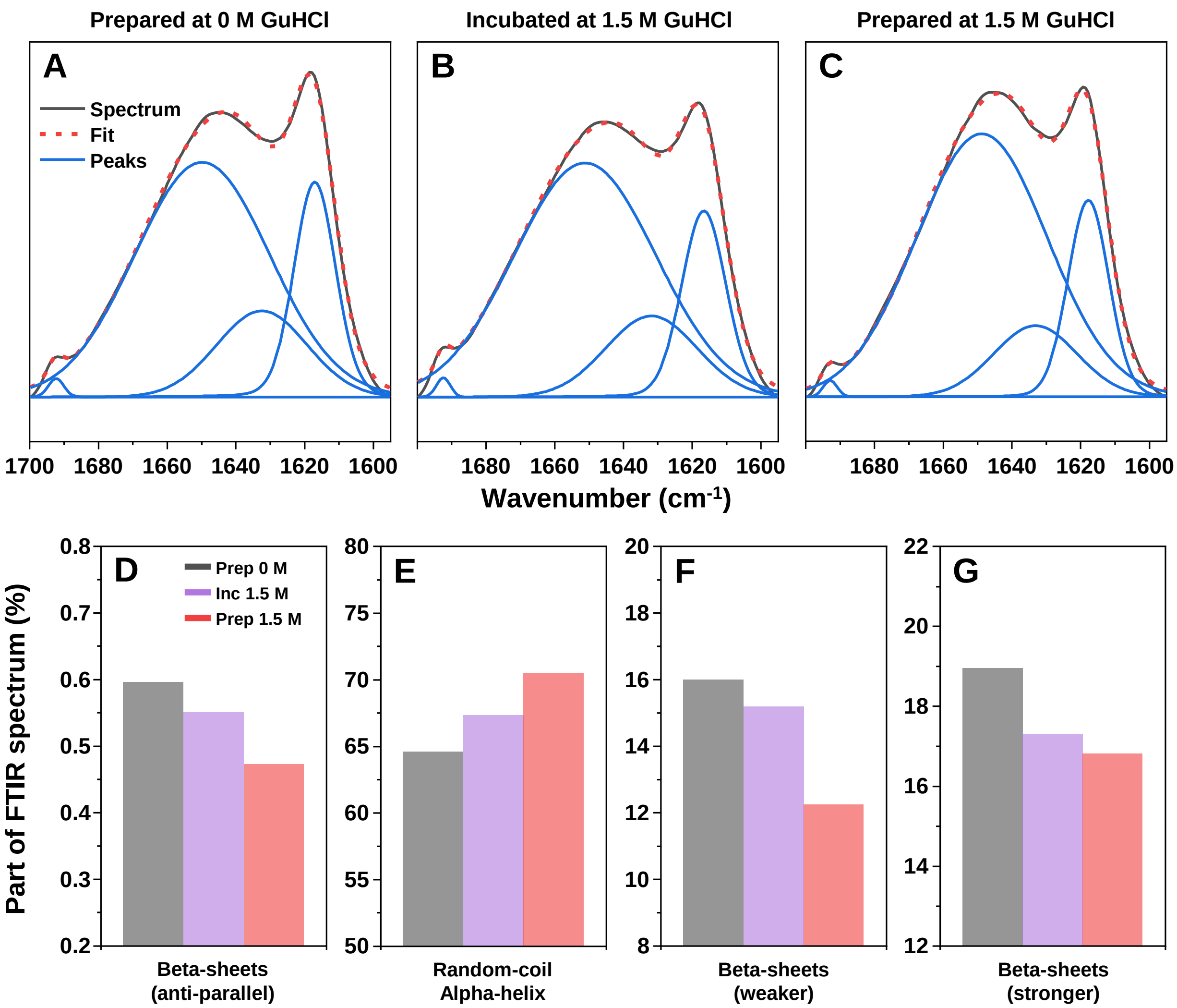


**Figure S3.** Deconvolution of S100A9 fibril FTIR spectra when they were formed under 0 M GuHCl conditions (A), resuspended and incubated under 1.5 M GuHCl (B) or formed under 1.5 M GuHCl conditions (C). The parts of FTIR spectra associated with anti-parallel beta-sheets (D), random-coil or alpha-helical motifs (E), weaker (F) and stronger (G) hydrogen bond strength parallel beta-sheets.

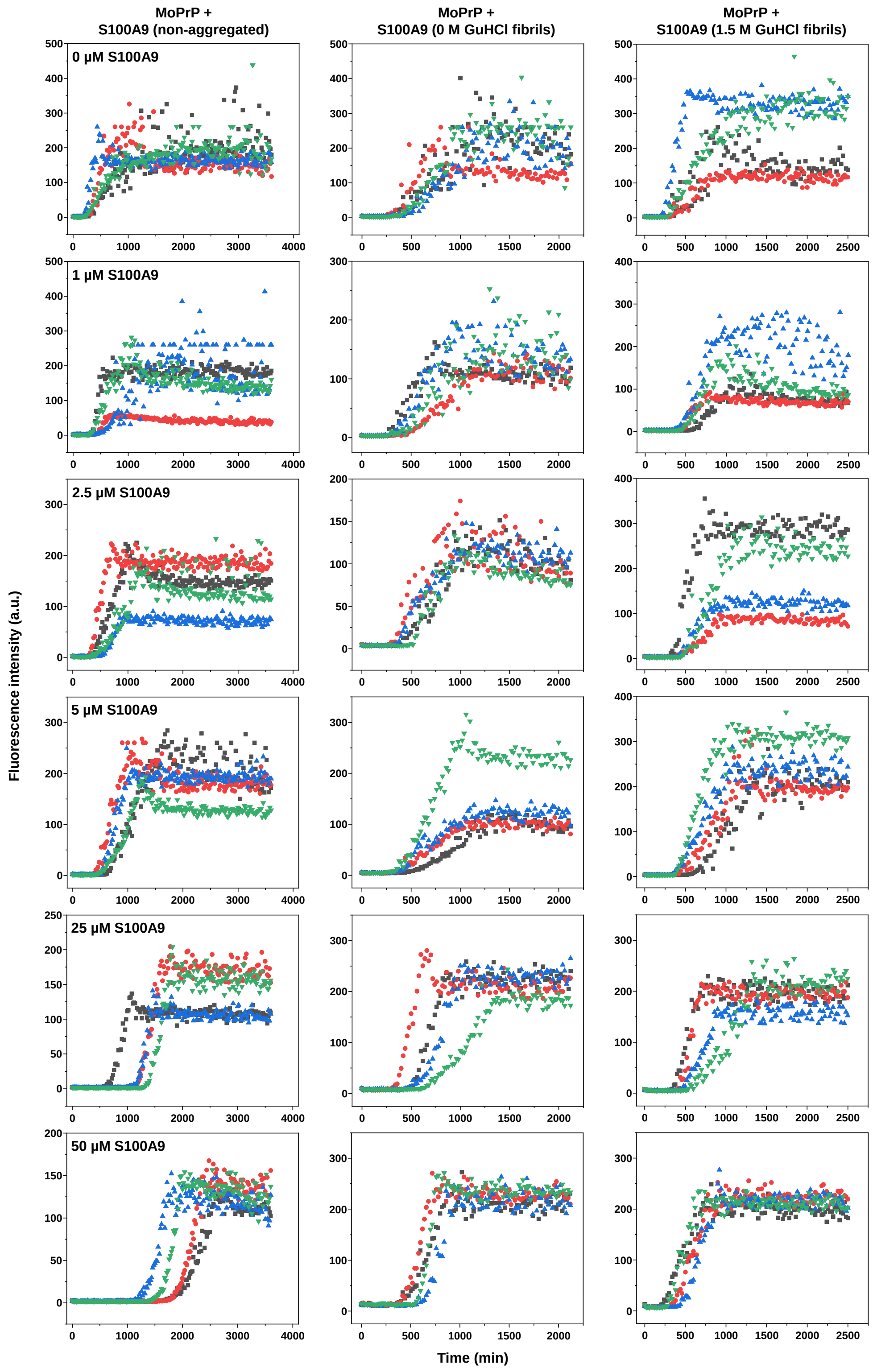


**Figure S4.** Representative aggregation curves (n=4) of PrP under different concentrations of non-aggregated S100A9 (first column) and S100A9 aggregated under 0 M GuHCl (second column) or 1.5 M GuHCl (third column) conditions.

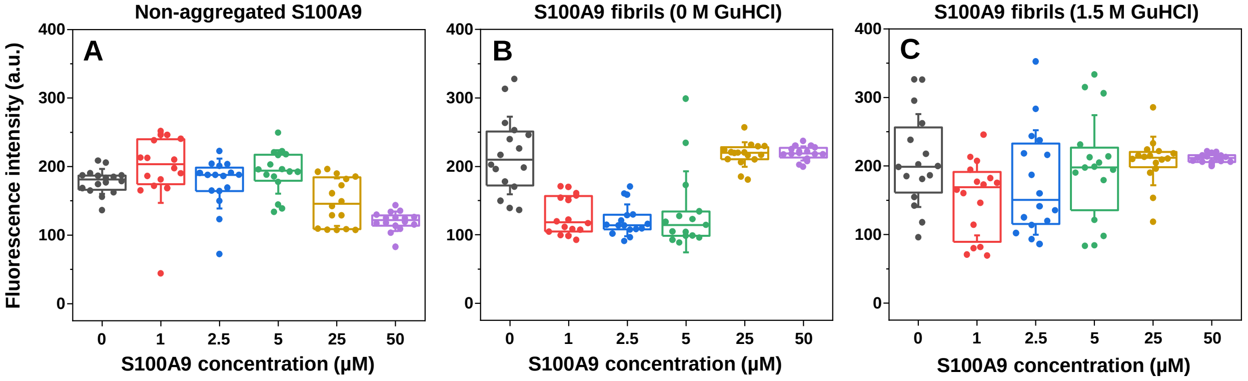


**Figure S5.** Fluorescence intensity distributions of PrP samples prepared under different conditions of non-aggregated S100A9 (A) and S100A9 fibrils prepared under 0 M (B) and 1.5 M GuHCl (C) conditions.

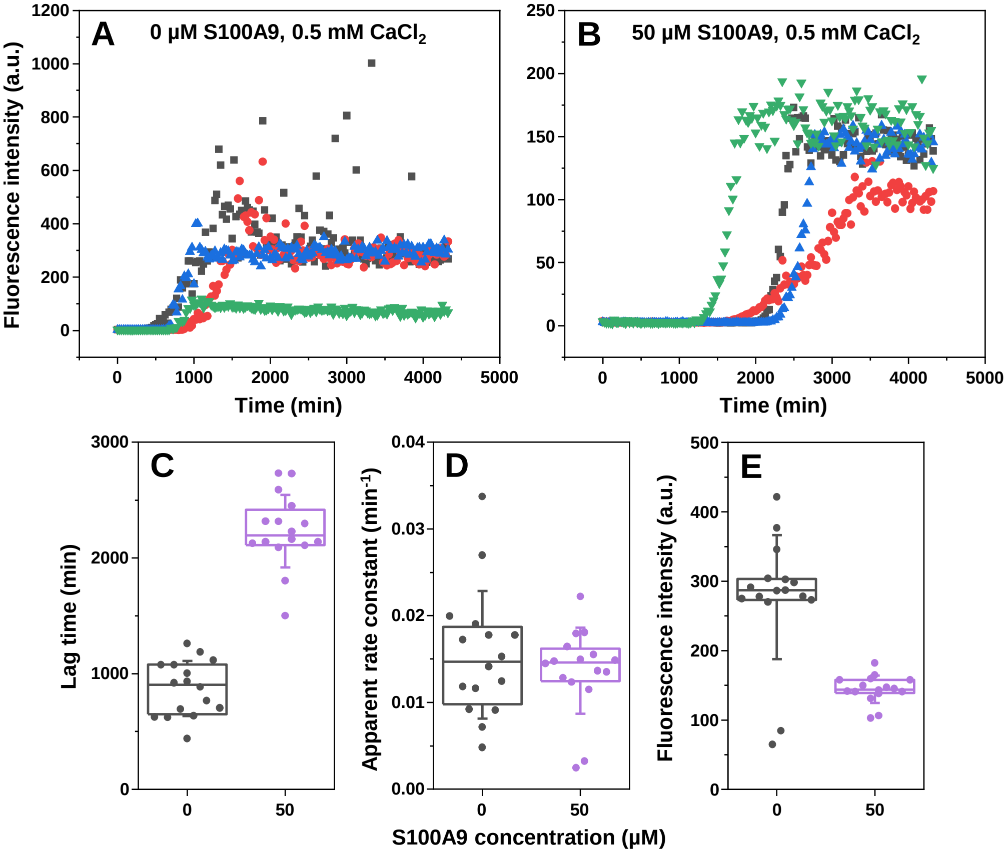


**Figure S6.** PrP aggregation in the presence of 0.5 mM $CaCl_2$ with and without 50 μM of non-aggregated S100A9. Representative aggregation curves (n=4) of PrP aggregation in the presence of 0.5 mM $CaCl_2$ and 0 μM (A) or 50 μM (B) non-aggregated S100A9. Lag time (C), apparent rate constant (D) and sample fluorescence intensity (E) distribution of both condition aggregation reactions (n=16, errors bars are for one standard deviation).

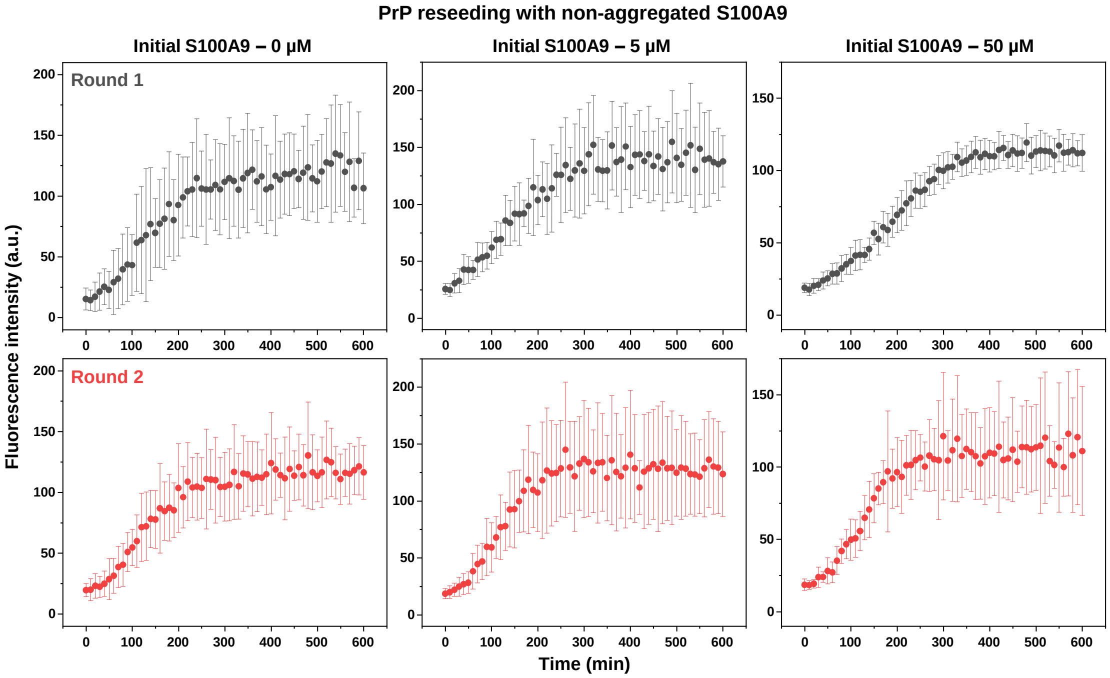


**Figure S7.** PrP reseeding aggregation curves of samples which initially contained different concentrations of non-aggregated S100A9. Each data point is the average of 16 repeats (one representative curve from each of the initial 16 sample reseeding procedures, error bars are for one standard deviation).

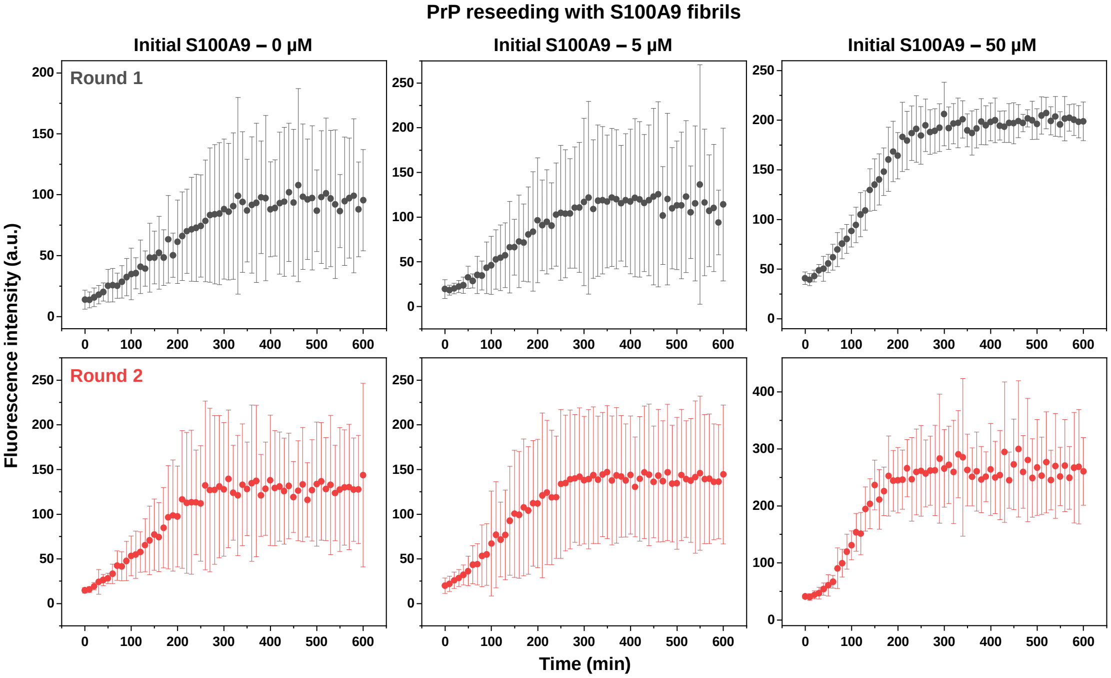


**Figure S8.** PrP reseeding aggregation curves of samples which initially contained different concentrations of S100A9 fibrils. Each data point is the average of 16 repeats (one representative curve from each of the initial 16 sample reseeding procedures, error bars are for one standard deviation).

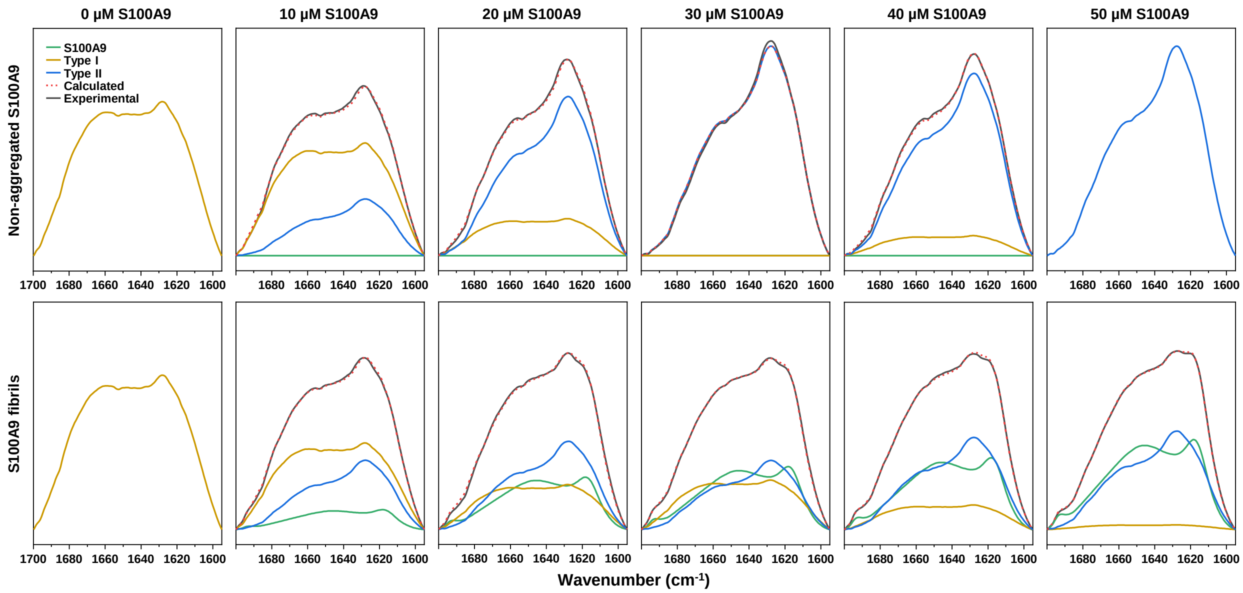


**Figure S9.** Deconvolution of PrP-S100A9 sample aggregate FTIR spectra. Part of FTIR spectra related to S100A9 fibrils (green), type I (yellow) and type II (blue) PrP fibrils. Combined S100A9, type I and II PrP fibril spectra (red dashed line) comparison with experimentally acquired S100A9-PrP spectra (black). Deconvolution results presented in Panels D and E were calculated from single bulk solution FTIR spectra of 6 combined samples for each condition.

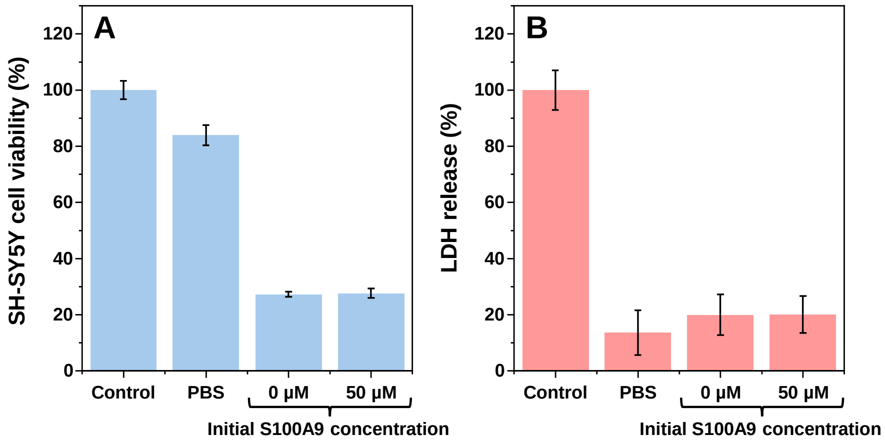


**Figure S10.** Effect of PrP aggregates on SH-SY5Y human neuroblastoma cells. Reseeded PrP aggregate, initially prepared with 0 µM or 50 µM non-aggregated S100A9, effect on cell metabolic activity (A, MTT assay) and their cytotoxicity (B, LDH assay). For each condition, three independent assays were carried out (each with three sample repeats), error bars are for one standard deviation (n=9).

### S100A9 cloning, expression and purification

Plasmid encoding His6-SUMO-S100A9 gene was derived from constructs of S100A9 (a gift of Ludmilla Morozova-Roche (Umeå)) and pET28_SUMO_PDZ1PDZ2 (a gift of Björn M. Burmann (Gothenburg)]. Briefly, the S100A9 gene and the His6-SUMO tag were amplified and fused by standard PCR methods, yielding the His6-SUMO-S100A9 insert which was cloned into the pET28a(-) vector (5'-NcoI; 3'-BamHI). All primers used for cloning are listed in Table 1.

**Table 1.** Plasmid, primers and primer sequences used for S100A9 cloning.

| Plasmid | Primer | Sequence |
|---|---|---|
| pDS99 (6xHis-SUMO-S100A9 | pet15_5206 | 5' ATCGAGATCTCGATCCCGCG 3' |
| | DS99_S100A9_BamHI | 5' CGGGATCCTTAGGGGGTGCCCTCCCC 3' |
| | DS99_S100A9_frw | 5' GATTGGCGGTATGACTTGCAAAATGTCGCAG 3' |
| | DS99_S100A9_rev | 5' GCAAGTCATACCGCCAATCTGTTCCAGATG 3' |

The plasmid containing His6-SUMO-S100A9 was chemically transformed into One ShotTM E. coli BL21 StarTM (DE3) (Fisher Scientific) cells, that afterwards were grown at 37°C in LB medium containing kanamycin (50 mg/ml). Once optical density was reached at 600 nm ≈ 0.7, protein production was induced with 0.4 mM isopropyl-β-d-thiogalactopyranoside (Fisher Scientific), and the cells were left to grow overnight at 25°C. Cells were harvested by centrifugation at 6000 x g for 20 min at 4°C and subsequently resuspended in 50 ml of lysis buffer (25 mM Hepes/NaOH, 0.5 M NaCl, and 10 mM imidazole (pH 7.5)). The suspension was disrupted with a Sonopuls (Bandelin) homogeniser (10 s on, 30 s off, 30% power, total time 30 min). Cell debris was removed by centrifugation at 18,000 x g for 45 min at 4°C, and the supernatant was applied to a $Ni^{2+}$ Sepharose 6 Fast Flow (Cytiva) loaded gravity column, followed by stepwise elution with 20 ml of lysis buffer supplemented with 75 and 300 mM imidazole, respectively. Fractions containing the His6-SUMO-S100A9 protein were dialyzed two times against 10 mM Tris buffer (pH 7.4) and the His6-SUMO tag was cleaved with human Sentrin-specific protease 1 (SENP1) catalytic domain (derived from pET28a-HsSENP1, that was a gift from Jorge Eduardo Azevedo (Addgene plasmid #71465) at 4°C overnight [1,2]. The cleaved proteins were applied again to a $Ni^{2+}$ column, and the flow-through was collected. The proteins were concentrated using Amicon centrifugal filters [10k Molecular weight cut-off (MWCO), Merck Millipore] and purified further by size exclusion chromatography (HiLoad® 26/600 Superdex® 75 pg, Cytiva) in PBS.

### References


1. Panavas, T., Sanders, C. & Butt, T. R. SUMO Protocols: Chapter 20. *Methods Mol. Biol.* **497**, 303–317 (2009).
2. Mendes, A. V., Grou, C. P., Azevedo, J. E. & Pinto, M. P. Evaluation of the activity and substrate specificity of the human SENP family of SUMO proteases. *Biochim. Biophys. Acta - Mol. Cell Res.* **1863**, 139–147 (2016).
